# Actigraphy in brain-injured patients – A valid measurement for assessing circadian rhythms?

**DOI:** 10.1101/839472

**Authors:** Monika Angerer, Manuel Schabus, Marion Raml, Gerald Pichler, Alexander B. Kunz, Monika Scarpatetti, Eugen Trinka, Christine Blume

## Abstract

**Background:** Actigraphy has received increasing attention in classifying rest-activity cycles. However, in patients with disorders of consciousness (DOC), actigraphy data may be considerably confounded by passive movements, such as nursing activities and therapies. Consequently, this study verified whether circadian rhythmicity is (still) visible in actigraphy data from patients with DOC after correcting for passive movements.

**Methods:** Wrist actigraphy was recorded over 7-8 consecutive days in patients with DOC (diagnosed with unresponsive wakefulness syndrome [UWS; *n*=19] and [exit] minimally conscious state [MCS/EMCS; *n*=11]). Presence and actions of clinical and research staff as well as visitors were indicated using a tablet in the patient’s room. Following removal and interpolation of passive movements, non-parametric rank-based tests were computed to identify differences between circadian parameters of uncorrected and corrected actigraphy data.

**Results:** Uncorrected actigraphy data overestimated the *inter*daily stability and *intra*daily variability of patients’ activity and underestimated the deviation from a circadian 24h rhythm. Only 5/30 (17%) patients deviated more than 1h from 24h in the uncorrected data, whereas this was the case for 17/30 (57%) patients in the corrected data. When contrasting diagnoses based on the corrected dataset, stronger circadian rhythms and higher activity levels were observed in MCS/EMCS as compared to UWS patients. Day-to-night differences in activity were evident for both patient groups.

**Conclusion:** Our findings indicate that uncorrected actigraphy data overestimates the circadian rhythmicity of patients’ activity, as nursing activities, therapies, and visits by relatives follow a circadian pattern itself. Therefore, we suggest correcting actigraphy data from patients with reduced mobility.

## Background

In the last decades, the measurement of physical activity, so-called actigraphy, has received increasing attention for the classification of vigilance states in healthy individuals (see reference 1 for a review). Recently, actigraphy was also used for the investigation of day/night patterns as well as circadian rhythms (i.e. rhythms with a period length of approximately 24 h) in patients following severe brain injury[2-5]. As those patients often need full-time care, actigraphy measures may be highly influenced by passive movements in this patient population. Therefore, we sought to systematically control for passive movements in this study.

Severe brain injury can cause coma and, upon recovery, longer lasting changes in consciousness, which can be summarized as “disorders of consciousness (DOC)”. In a simplified approach, consciousness is thought to require both adequate levels of wakefulness and awareness[6]. More precisely, wakefulness refers to some degree of arousal at brain level (e.g. eye-opening, limb movements) and awareness denotes the ability to have a conscious experience of any kind. While brain-dead or comatose patients are characterized by absent arousal and awareness, patients with an unresponsive wakefulness syndrome (UWS; formerly often referred to as vegetative state) show some return of arousal (i.e. alternating phases of sleep [closed eyes] and wakefulness [opened eyes]), however, without signs of awareness. In a minimally conscious state (MCS), cognitively mediated behavior indicating awareness occurs inconsistently, but is reproducible or long enough to be differentiated from reflexive behavior (e.g. response to command, verbalizations, visual pursuit)[7]. If patients are able to functionally use objects and communicate, their state is denoted exit MCS (EMCS)[8]. Thus, while UWS patients are assumed to be unconscious, MCS and EMCS patients show signs of consciousness. However, distinguishing between UWS and MCS is still a challenging task. Until now, behavioral methods like the “Coma Recovery Scale – Revised” (CRS-R)[9] and the “Glasgow Coma Scale” (GCS)[10] remain the best available tools for clinical diagnoses. Unfortunately, the rate of misdiagnoses is still high (∼40%)[11] if behavioral scales are not performed by well-trained professionals. Therefore, the quest for ways to improve the validity of such assessments remains an important issue. As consolidated periods of wakefulness and sleep resulting from well-entrained circadian rhythms, seem crucial for adequate arousal levels and thus (conscious) wakefulness, circadian rhythms have been the focus of recent research in patients with DOC. Research from our group[5, 12] suggests that a better integrity of patients’ circadian melatonin(-sulfate) and temperature rhythms is indeed related to a richer behavioral repertoire (as measured with the CRS-R). Knowing a patient’s circadian rhythm in turn has been suggested to help find the optimal time for behavioral assessments and therapies as cognitive functions also vary with the time of day[12-14]. However, besides temperature and melatonin rhythms, variability within a day can also be observed in other parameters in patients with DOC as for example in blood pressure, heart rate and body movements[3, 15, 16].

Body movements can be monitored through actigraphy, which is frequently used in the clinical setting for evaluating rest-activity cycles (e.g. in insomnia, circadian rhythm disorders, or clinical monitoring in the rehabilitation process of patients with traumatic brain injury (TBI) [17, 18]) with the major advantage of being a cost-efficient and easy to use tool suitable for long-term investigations. More precisely, an actigraph, worn on the wrist or ankle allows the continuous recording of data across days, weeks and even months in a natural setting without restricting mobility and daily life routine of the participants.

Previous studies investigating rest-activity cycles in patients with DOC using actigraphy found that the sleep-wake cycle deteriorates with decreasing consciousness level[2]. When taking etiology into account, only patients with TBI show significant day-night differences (i.e. stronger motor activity during day time [7 am – 11 pm] than during night-time [11 pm – 7 am]) but not patients with anoxic-ischemic brain injuries (AI)[4]. Furthermore, circadian sleep-wake cycles (that is, not only day-to-night variations but the investigation of fluctuations in wrist actigraphy-derived physical activity over several days using cosinor rhythmometry analyses) are more impaired in UWS patients and patients with non-traumatic brain injuries (NTBI) as compared to MCS patients and patients with TBI. Therefore, Cruse et al. suggest that actigraphy should be considered as an alternative for assessing sleep-wake cycles in patients with DOC and appeal to also determine the prognostic utility of wrist actigraphy for UWS and MCS patients in future studies[3].

However, the use of actigraphy in patients with DOC may be severely limited by several factors. First, patients with DOC often suffer from severe motor impairments, spasticity and the use of muscle relaxants, which have previously shown to, descriptively, decrease for example the concordance between polysomnography and actigraphy-derived parameters[18]. Second, as most of them spend much time in bed and often need full-time care in hospitals or nursing homes, actigraphy data is likely to be confounded by passive movements due to nursing activities, therapies or movements initiated by visitors. The latter issue becomes particularly crucial when actigraphy data are used to make inferences about patients’ circadian rhythms. This is because the rhythmicity might reflect daily patterns of e.g. nursing activities or therapies rather than a circadian rhythm of the patient. Unfortunately, correcting for passive movements is challenging and the previously published findings may thus be biased towards overestimating rhythmicity. In the current paper, we therefore sought to systematically control for passive movements and to assess the magnitude of the introduced bias by comparing corrected and uncorrected actigraphy-derived measures. Eventually, we aimed at revealing whether circadian rhythmicity can be identified in MCS and/or UWS patients using actigraphy data even if artificial biases are carefully controlled for.

## Methods

### Patients

From a total of 30 patients one patient (P26) had to be excluded because hardly any activity was left after cleaning the data from passive movements (cf. *Additional File 1: Tables S1, S2* and *Figure S2*). Thus 29 patients (13 women) aged 19-78 (mdn = 55 years) from long-term care facilities in Austria were included in the study sample with 18 patients who were diagnosed with UWS (7 women), 7 were in a MCS (4 women) and 4 in an EMCS (2 women). Note that the data has been used in two previous publications, where we studied circadian rhythms in patients with DOC but without focusing on actigraphy data[5, 12]. Informed consent was obtained from the patients’ legal representatives and the study had been approved by the local ethics committees. Please note that MCS and EMCS patients were combined to a single group in the analyses as we sought to analyze differences between unconscious UWS and (minimally) conscious (E)MCS patients. For more details on the study sample please see *Table 1*.

**Table 1.** Demographic information.

| Patient ID | Age | Gender | Etiology | Time since injury (months) | Diagnosis | CRS-R sum score |
| --- | --- | --- | --- | --- | --- | --- |
| P1 | 43 | M | NTBI | 39.0 | EMCS | 11 |
| P2 | 72 | F | NTBI | 10.0 | UWS | 6 |
| P3 | 25 | M | NTBI | 99.0 | UWS | 6 |
| P4 | 34 | M | TBI | 15.0 | UWS | 6 |
| P5 | 60 | M | NTBI | 7.0 | UWS | 7 |
| P6 | 49 | F | NTBI | 16.0 | UWS | 6 |
| P7 | 50 | M | NTBI | 4.0 | UWS | 3 |
| P8 | 59 | F | NTBI | 6.0 | UWS | 6 |
| P9 | 60 | M | NTBI | 7.0 | UWS | 7 |
| P10 | 68 | M | NTBI | 5.0 | UWS | 6 |
| P11 | 70 | F | TBI | 7.0 | UWS | 3 |
| P12 | 48 | F | NTBI | 37.0 | UWS | 5 |
| P13 | 66 | M | NTBI | 2.0 | UWS | 7 |
| P14 | 20 | M | TBI | 56.0 | MCS | 13 |
| P15 | 71 | M | NTBI | 24.0 | UWS | 1 |
| P16 | 55 | F | TBI | 168.0 | MCS | 17 |
| P17 | 70 | F | NTBI | 15.0 | UWS | 3 |
| P18 | 51 | M | TBI | 54.0 | UWS | 4 |
| P19 | 61 | F | NTBI | 9.0 | EMCS | 23 |
| P20 | 68 | M | NTBI | 415.0 | UWS | 4 |
| P21 | 53 | F | NTBI | 10.5 | MCS | 13 |
| P22 | 68 | F | TBI | 13.5 | MCS | 9 |
| P23 | 71 | F | TBI | 2.5 | EMCS | 23 |
| P24 | 53 | F | NTBI | 82.0 | UWS | 5 |
| P25 | 37 | M | TBI | 197.0 | MCS | 9 |
| P26 | 46 | F | NTBI | 3.0 | UWS | 4 |
| P27 | 19 | F | TBI | 17.0 | MCS | 8 |
| P28 | 78 | M | NTBI | 13.0 | MCS | 9 |
| P29 | 27 | M | NTBI | 1.5 | UWS | 4 |
| P30 | 54 | M | TBI | 10.0 | EMCS | 20 |

M = male; F = female; NTBI = non-traumatic brain injury; TBI = traumatic brain injury; UWS = unresponsive wakefulness syndrome; MCS = minimally conscious state; EMCS = Exit MCS; CRS-R = Coma Recovery Scale – Revised.

### Experimental Design

The study protocol comprised seven to eight full days (hereinafter “study week”) during which actigraphy was assessed continuously (for further measures recorded see reference 5). Patients’ behavioral repertoire or level of consciousness was assessed with the CRS-R in the morning of day 6 and in the afternoon of day 7 during the study week. Besides this, multiple additional CRS-R assessments (i.e. 10 additional assessments) were obtained in 16 patients (8 women; P2, P4, P6, P8, P10, P12, P14, P16, P18; P24-P30) on two consecutive days following the study week (note that multiple CRS-R assessments are not available for all patients as they were added to the study protocol later). Illuminance was kept <500 lux at eye level during the day (7 am – 9 pm) and <10 lux during the night (9 pm – 7 am), which was ensured by continuous measurements with light sensors (wGT3X-BT Monitor, ActiGraph LLC., Pensacola, USA) and spot checks with a luxmeter (Dr. Meter, Digital Illuminance/Light Meter LX1330B). For further information on light levels please refer to *Additional File 1*.

### Behavioral Assessment and Data Analysis

#### Coma Recovery Scale – Revised

The patients’ neurophysiological state was assessed behaviorally with the CRS-R [9]. It is composed out of six subscales reflecting auditory, visual, motor, oromotor, communication and arousal functions that altogether make up 23 items. Whereas the lowest item on each subscale represents reflexive behavior, the highest item indicates cognitively mediated behavior. Patients are tested in a hierarchical manner; meaning that the examiner starts with the highest item of each subscale and moves down the scale until the patient’s response meets the criteria for one item. The scores of all subscales sum up to a maximum score of 23. The assessment was done twice by two trained experts in all patients, with 10 additional assessments being available for 16 patients. For the following analyses we used those CRS-R assessments where the patients showed the highest behavioral reactivity (e.g. as characterized by the best diagnosis or highest sum score) as this is thought to best represent the true state of the patient. The highest CRS-R score and diagnosis across the whole study period of each patient are shown in *Table 1.* For further information on multiple CRS-R assessments please refer to *Additional File 1.*

#### Actigraphy

We recorded actigraphy with a sampling rate of 30 Hz using GT3X+ devices (ActiGraph LLC., Pensacola, FL 32502). The actigraph was placed on the wrist of the arm with the greatest mobility and least spasticity. If both arms were equally mobile, it was placed on the wrist of the dominant hand. If the legs were more mobile it was placed on the ankle of the most mobile leg. Actigraphs recorded continuously during the whole “study week” and were only taken off if the patients were showered or bathed. To monitor passive movements and remove artifacts resulting from them, we recorded all events deemed relevant in the patient room using an application (https://github.com/wolli2710/HospitalTracker) that enabled clinical and research staff as well as visitors to indicate the type of activity that was performed by simply tapping the screen of a tablet in the patient room. Specifically, we had start and end buttons for visits, nursing activities, actigraphy (i.e. to mark if the actigraph was taken off for showering or bathing), therapy, mobilizations in the wheelchair and mobilizations outside the building (e.g. if they went for a walk with the patient). Furthermore, we had “single press buttons” (i.e. no start and stop option; only needed to be pressed once at the time of occurrence) for the administration of medication, nutrition as well as for lights on and out, eyes open and closed (cf. *Additional File 1: Figure S1* to get an impression of the graphical user interface of the tablet). Upon tapping the screen, a time stamp was generated, which allowed us to correct the actigraphy data post hoc.

Cleaning and analysis of actigraphy data was done in R version 3.4.2[19]. After integrating actigraphy and tablet data into one single dataset, the actigraphy data was down-sampled to 1/60 Hz (i.e. one value per minute). The actigraphy values of the time spans during which (i) clinical staff or visitors were with the patient, (ii) the patient was put into a wheelchair or back into bed, (iii) the CRS-R assessments took place as well as (iv) the times when the actigraphs had been taken off for body care, were removed. As the calculation of interdaily stability (IS; see below) requires a dataset without missing data, the first half of the removed values was replaced by the median activity during the 10 min preceding the event and the second half was replaced by the median activity during the 10 min following the end of the event. Importantly, to account for the issue that clinical staff or visitors indicated their presence too late, we additionally removed and imputed 5 min before and after each nursing activity as well as 10 min before and after each visit or usage of the wheelchair. This automatic artefact correction was followed by a visual screening and manual correction of residual artefacts. Thus, the resulting dataset can be assumed to be free from passive movements representing only the “true” internal motor activity of the patient (cf. *Figure 1* for an illustration of our correction procedure). For the analyses of the uncorrected actigraphy data we down-sampled the data to 1/60 Hz. Thus, we arrived at a corrected as well as at an uncorrected dataset for each patient, which we used for the calculation of the following parameters using R.

**Figure 1.**
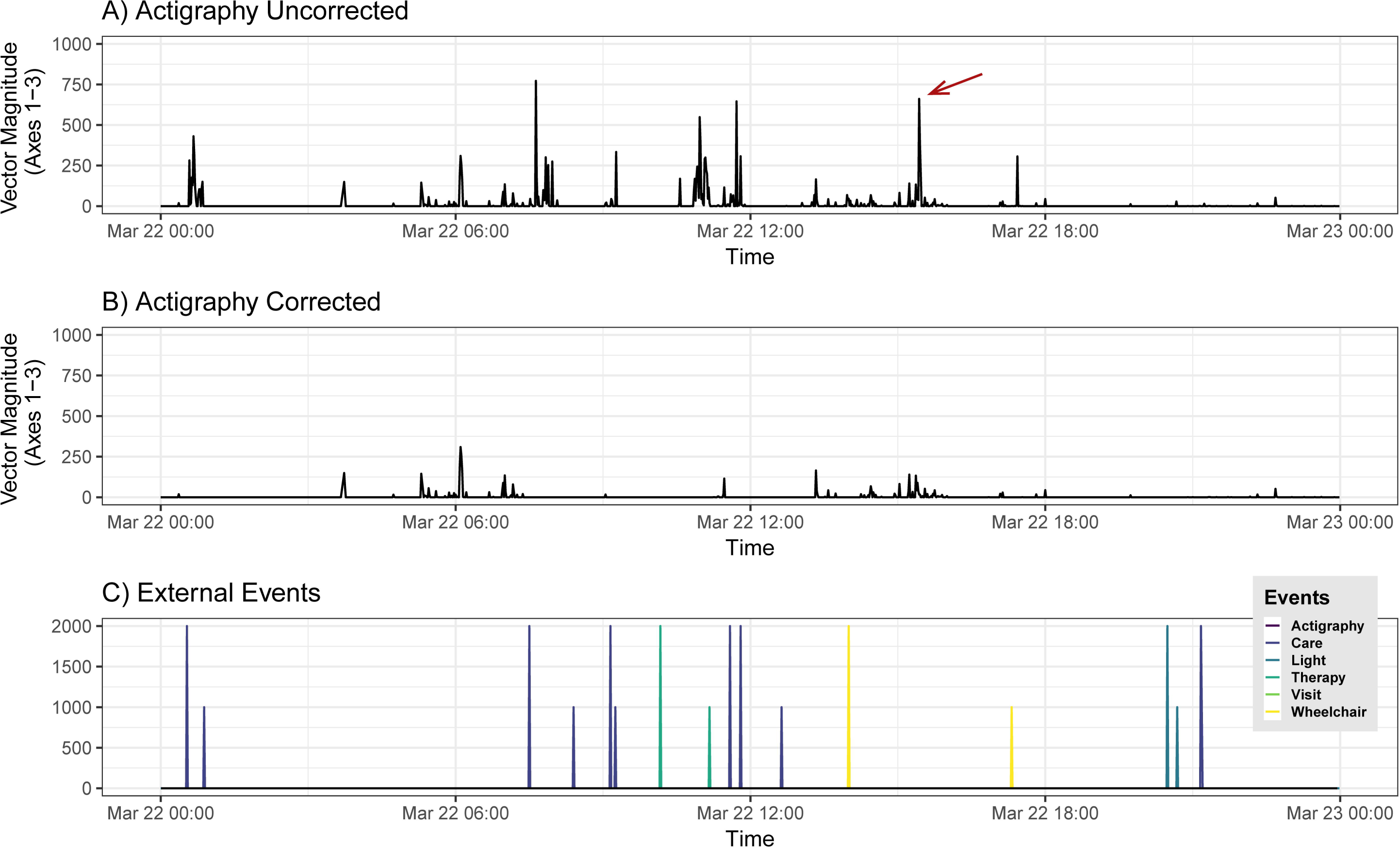
Graphical representation of the manual and automatic artefact correction of a 24 h actigraphy recording. A) Uncorrected actigraphy data with the time of day being depicted on the x-axis and the amplitude of the motor activity on the y-axis. B) Corrected actigraphy data after automatic (according to the tablet data) and manual artefact correction (marked with a red arrow). C) External events recorded by the tablet in the patient room with longer vertical lines representing the start and shorter vertical lines the stop of the respective event.

#### Interdailiy Stability and Intradaily Variability

Interdaily stability (IS) and intradaily variability (IV) are non-parametric measures[20], whose calculation is implemented in the R package ‘nparACT’[21]. In more detail, IS reflects how well a patient’s activity rhythm is entrained to a 24 h zeitgeber (i.e. the light-dark cycle) as indexed by values ranging between 0 for Gaussian noise and 1 for perfect IS. In contrast, IV quantifies the fragmentation of a rest-activity pattern. IV converges to 0 for a perfect sine wave and approaches 2 for Gaussian noise. It may even be higher than 2 if a definite ultradian component with a period length of 2 h is present in the rest-activity cycle. For individual patients’ results please refer to *Additional File 1: Tables S1* and *S2*.

#### Lomb-Scargle Periodograms

To detect rhythmicity in our data, we computed Lomb-Scargle periodograms[22, 23]. For each patient, we calculated two parameters using the “lomb” package available for R[24]: (1) normalized power and (2) peak period. The normalized power describes the fit of a sine wave to the data. It is maximal where the sum of squares of the fitted sine wave to the data is minimal. For calculation of the period length of each patient’s activity rhythm, we looked for significant peaks in the normalized power of the periodogram and extracted the period length of the significant peak, which was closest to 24 h (i.e. as circadian rhythms should be entrained to a 24 h cycle in a natural setting which is close to the intrinsic period of the human circadian pacemaker that is on average 24.18 h[25]). We set the oversampling factor to 100 and the significance level to *α* = 0.001. The individual patients’ results are displayed in *Additional File 1: Tables S1* and *S2*. For further information on the analyses please refer to the supplementary material of Blume et al.[5]

#### Mean Activity

Mean Activity was calculated separately for day (7 am – 9 pm) and night-time (9 pm – 7 am) and simply reflects the mean of the measured activity during the study week (arbitrary units). It takes the intensity and number of movements into account. For individual patients’ results please refer to *Additional File 1: Tables S1* and *S2*.

### Statistical Analyses

Statistical analyses were done in R. We investigated differences in actigraphy (IS, IV, normalized power, deviation of the peak period from a 24 h rhythm, mean activity) between corrected and uncorrected data as well as day-night differences in mean activity using Wilcoxon signed-rank test. Differences between diagnoses (i.e., UWS vs. MCS/EMCS) and etiology (i.e., TBI vs NTBI) were investigated using Mann-Whitney U test. To check if the differences in actigraphy data between UWS and MCS/EMCS patients are also visible on a subscale level, we also investigated the correlation between patients’ CRS-R scores (sum score as well as subscale scores) and actigraphy data using Kendall’s Tau. The significance level was *α* = .05 (two-sided) with *p*-values .05 < *p* ≤ .1 being denoted trends. Regarding effect sizes, 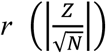 was calculated for the results of Wilcoxon signed-rank test and Mann-Whitney U test. According to Cohen[26], the following conventions are applied when interpreting *r*: small effect: *r* = .1; medium effect: *r* = .3; large effect: *r* = .5.

## Results

### Circadian Rhythms

Comparisons between corrected and uncorrected actigraphy data revealed that interdaily stability (IS) (*Z*(*N=29*)=-2.96, *p*=.003, *r*=.55; cf. *Figure 2 A*) and IV (*Z*(*N=29*)=-4.22, *p*<.001, *r*=.78; cf. *Figure 2 B*) were higher in the uncorrected data than in the corrected data. The period length was closer to 24 h in the uncorrected data (*Z*(*N=29*)=-3.29, *p*=.001, *r*=.61; median deviation from 24 h: uncorrected data=0.41 h, corrected data=1.11 h; cf. *Figure 3 A*). The strength of the circadian rhythm (i.e. normalized power) did not differ between datasets (*Z*(*N=29*)=-.86, *p*=.39, *r*=.16; cf. *Additional File 1: Figure S3*).

**Figure 2.**
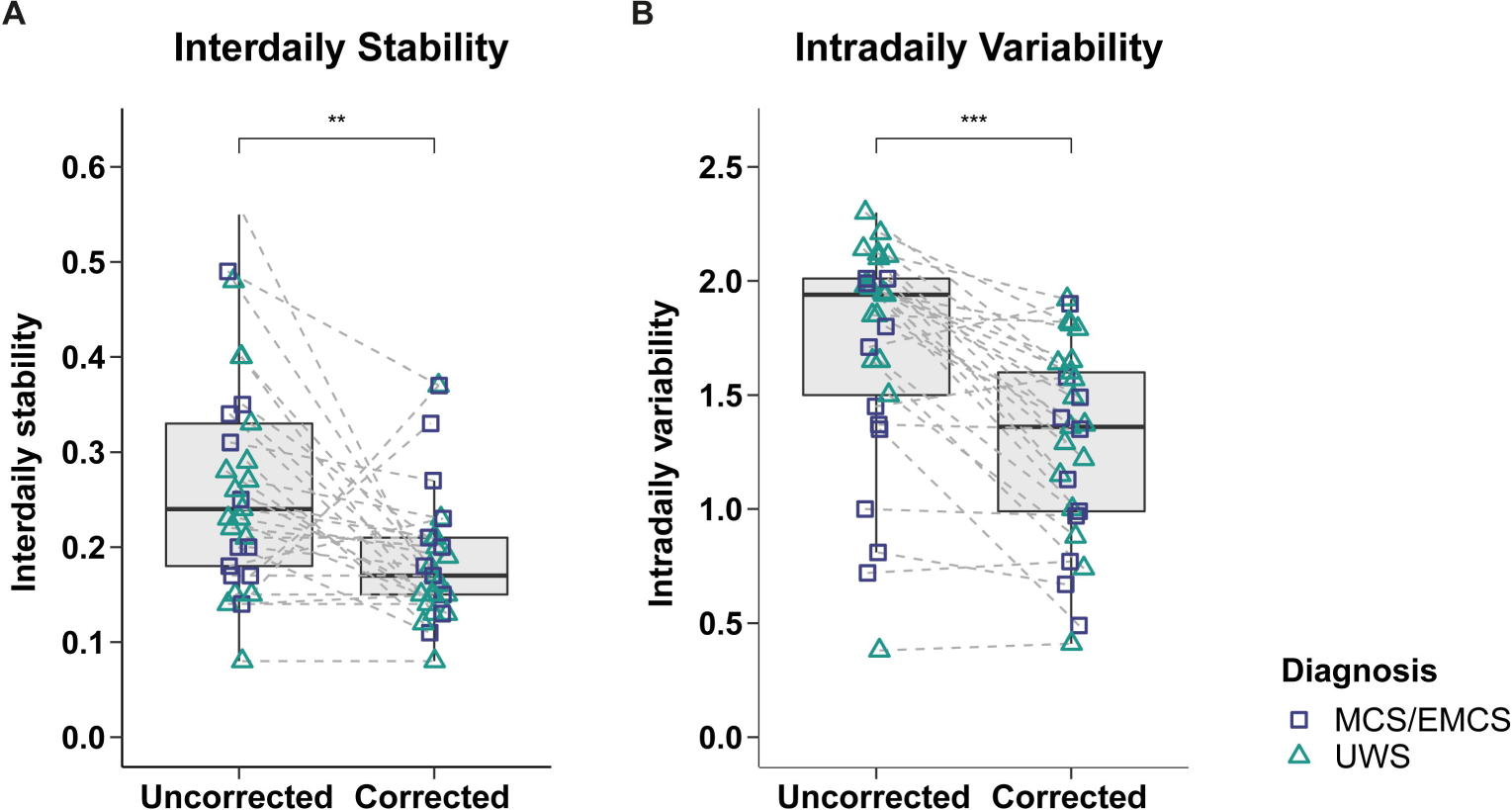
Interdaily stability (A) and intradaily variability (B) in uncorrected vs. corrected data. **A) Interdaily stability (IS).** The IS was overestimated and significantly higher in the uncorrected data (IS approaches 0 for Gaussian noise and converges to 1 for perfect IS.). UWS and MCS/EMCS patients did not differ in both corrected and uncorrected data *(cf. Additional File 1: Figures S4 A-B)*. **B) Intradaily variability (IV).** The IV was also overestimated and significantly higher in the uncorrected data (IV converges to 0 for a perfect sine wave [i.e. no IV] and approaches 2 for Gaussian noise. Values > 2 indicate an ultradian component with a period length of 2 h.). UWS and MCS/EMCS patients only differed in the uncorrected data *(cf. Additional File 1: Figures S4 C-D)*. Horizontal lines represent the medians, boxes the interquartile range (IQR; distance between the 1^st^ [Q1] and 3^rd^ quartile [Q3]), whiskers extend at most to Q1-1.5*IQR (lower whisker) and Q3+1.5*IQR (upper whisker). Asterisks indicate significance: \*\*\**p* ≤ .001, \*\**p* ≤ .01. Abbreviations: MCS = minimally conscious state, EMCS = Exit MCS, UWS = unresponsive wakefulness syndrome.

**Figure 3.**
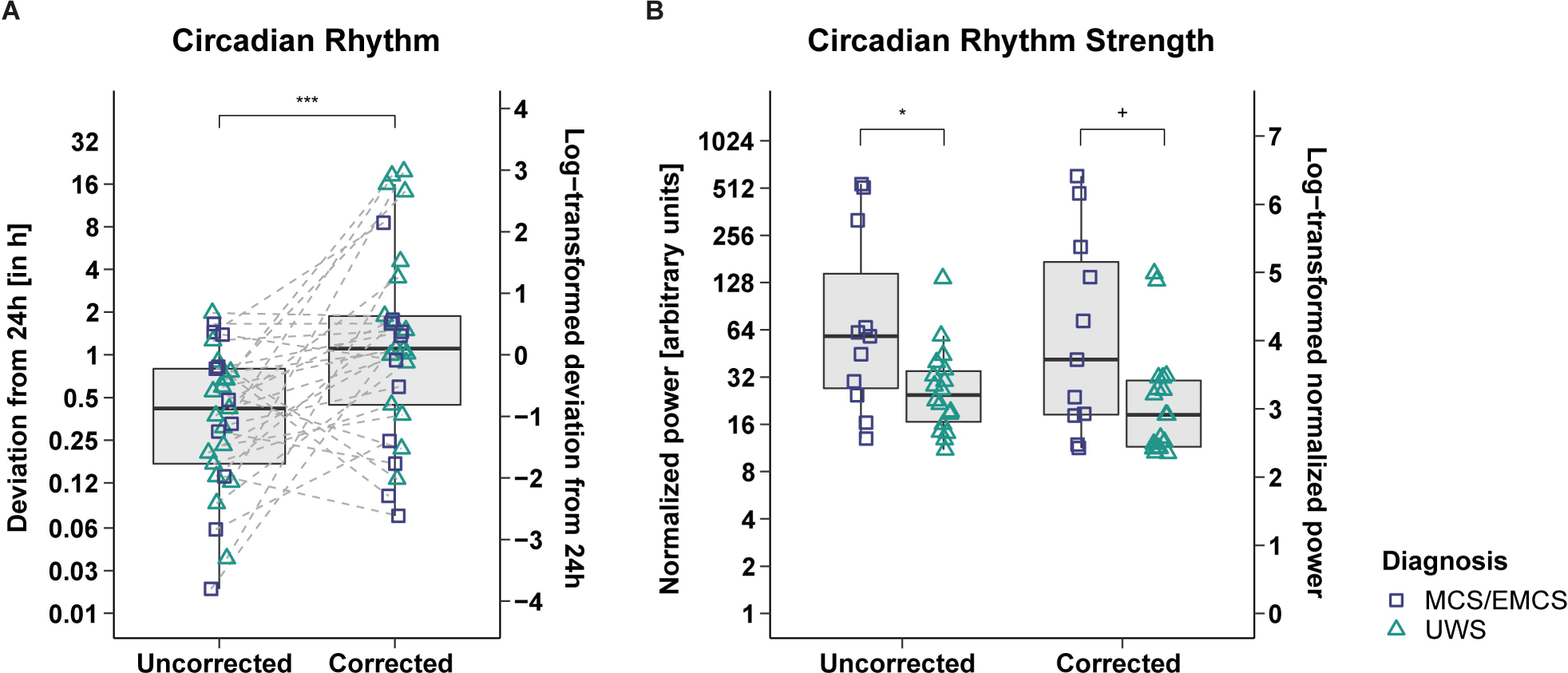
Circadian rhythmicity contrasted between datasets (A) and circadian rhythm strength contrasted between diagnoses (B). **A) Deviation of the patients’ peak period from 24 h.** The patients’ activity rhythms were significantly better aligned with a 24 h-rhythm in the uncorrected data (=less deviation from 24 h). UWS and MCS/EMCS patients did not differ in both uncorrected and corrected data *(cf. Additional File 1: Figures S4 E-F)*. **B) Normalized power of the patients’ peaks closest to 24 h.** UWS and MCS/EMCS patients differed in the uncorrected and corrected data. Pooling both patient groups the normalized power did not differ between datasets *(cf. Additional File 1: Figure S3)*. For better illustration, the data was log-transformed (right-hand y-axes); statistics were performed on the untransformed data (left-hand y-axes). Horizontal lines represent the medians, boxes the interquartile range (IQR; distance between the 1^st^ [Q1] and 3^rd^ quartile [Q3]), whiskers extend at most to Q1-1.5*IQR (lower whisker) and Q3+1.5*IQR (upper whisker). Asterisks indicate significance: \*\*\**p* ≤ .001, \**p* ≤ .05, ^+^p ≤ .1. Abbreviations: MCS = minimally conscious state, EMCS = Exit MCS, UWS = unresponsive wakefulness syndrome.

Contrasts between diagnoses showed that intradaily variability (IV) was higher in UWS patients than in MCS/EMCS patients in the uncorrected data (*Z*(*n1*=11, *n2*=18)=-2.20, *p*=.028, *r*=.41; cf. *Additional File 1: Figure S4 C*). This was not the case in the corrected data (*Z*(*n1*=11, *n2*=18)=-1.42, *p*=.157, *r*=.26; cf. *Additional File 1: Figure S4 D*). Furthermore, while MCS/EMCS patients showed a stronger circadian rhythm – as indicated by a higher normalized power – than UWS patients in the uncorrected data (*Z*(*n1*=11, *n2*=18)=2.16, *p*=.031, *r*=.40), this difference was only visible by trend in the corrected data (*Z*(*n1*=11, *n2*=18)=1.84, *p*=.065, *r*=.34; cf. *Figure 3 B*). Moreover, we found no significant differences between etiologies (TBI vs. NTBI) on any of the circadian rhythm indices in the corrected dataset (cf. *Additional File 1: Figures S5 A-D*).

### Day vs. Night

Patients’ activity levels were higher during day than night in both uncorrected (*Z*(*N=29*)=-4.13, *p*<.001, *r*=.77) and corrected data (*Z*(*N=29*)=-3.31, *p*<.001, *r*=.61) with effect sizes being larger in the uncorrected data (cf. *Additional File 1: Figure S6*). Furthermore, day-night differences were more pronounced in MCS/EMCS patients than in UWS patients in both datasets (uncorrected data: MCS/EMCS: *Z*(*n*=11)=-2.89, *p*=.004, *r*=.87; UWS: *Z*(*n=18*)=-2.92, *p*=.004, *r*=.69; corrected data: MCS/EMCS: *Z*(*n=11*)=-2.45, *p*=.014, *r*=.74; UWS: *Z*(*n=18*)=-2.22, *p*=.026, *r*=.52; cf. *Figure 4*).

**Figure 4.**
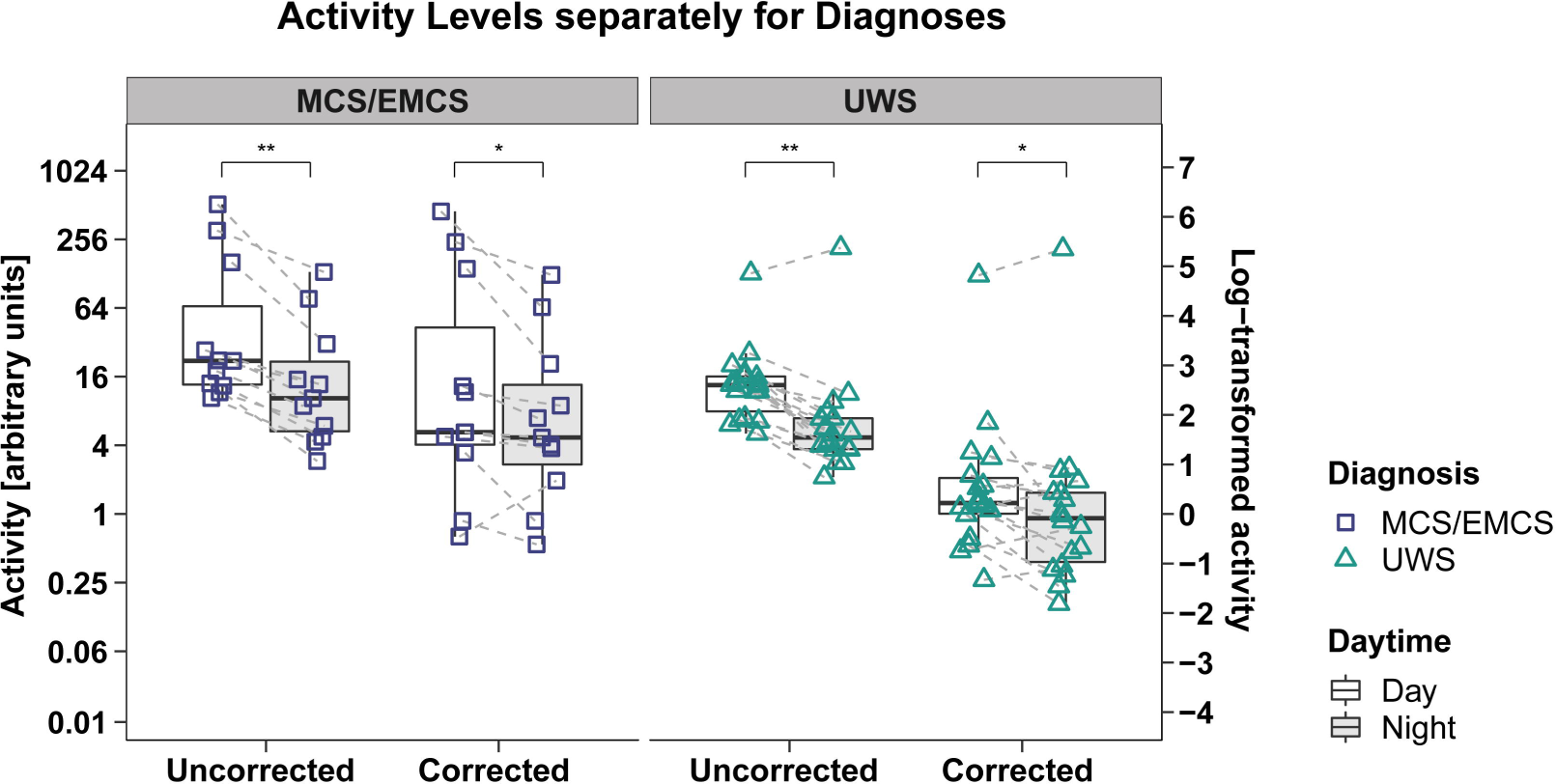
Patients’ mean activity during day vs. night in uncorrected and corrected data separately for diagnoses. The mean activity was significantly higher during the day (7am – 9pm) than during the night (9pm – 7am) in both uncorrected and corrected data in UWS and MCS/EMCS patients with stronger day-night effects in MCS/EMCS patients and uncorrected data. For better illustration, the data was log-transformed (right-hand y-axes); statistics were performed on the untransformed data (left-hand y-axes). Horizontal lines represent the medians, boxes the interquartile range (IQR; distance between the 1^st^ [Q1] and 3^rd^ quartile [Q3]), whiskers extend at most to Q1-1.5*IQR (lower whisker) and Q3+1.5*IQR (upper whisker). Asterisks indicate significance: \*\**p* ≤ .01, \**p* ≤ .05. Abbreviations: MCS = minimally conscious state, EMCS = Exit MCS, UWS = unresponsive wakefulness syndrome.

When comparing activity levels during day and night between diagnoses, we found that MCS/EMCS patients show higher mean activity than UWS patients during day and night in both uncorrected (day: *Z*(*n1*=11, *n2*=18)=2.16, *p*=.031, *r*=.40; night: *Z*(*n1*=11, *n2*=18)=2.20, *p*=.028, *r*=.41) and corrected data (day: *Z*(*n1*=11, *n2*=18)=-2.69, *p*=.007, *r*=.50; night: *Z*(*n1*=11, *n2*=18)=3.06, *p*=.002, *r*=.57) with larger effect sizes for comparisons between diagnoses in the corrected dataset (cf. *Additional File 1: Figures S7 A-D*). When looking at etiology, we found no significant differences between patients with TBI and NTBI in the activity levels during day and night in the corrected dataset (cf. *Additional File 1: Figure S8*).

## Discussion

Our results indicate that actigraphy data from clinical populations suffering from severe motor impairments such as patients with DOC is strongly influenced by passive movements, i.e. movements not initiated by the patients. Not correcting for these passive movements leads to an overestimation of the patients’ circadian rhythmicity rendering the validity of the uncorrected data highly questionable.

Analyses revealed that using uncorrected data resulted in an overestimation of how well patients’ circadian rhythms were entrained to a 24 h zeitgeber (as indicated by interdaily stability [IS] and the deviation from the peak closest to 24 h in the periodogram analyses) and in more pronounced day-night differences. Specifically, 25/30 patients (83%) showed a circadian rhythm (i.e. deviation less than 1 h from 24 h) in the uncorrected data (cf. *Additional File 1: Table S1*). This is well in line with the results from Cruse et al.[3] who found a circadian rhythm in 46/55 patients (84%). However, after correcting the actigraphy data for passive movements we found a circadian rhythm in only 13/30 patients (43%) (cf. *Additional File 1: Table S2*). This is most probably because nursing activities, therapies, and visiting times that cause such passive movements follow a regular (daily) schedule and are more prominent during the day than during the night. Thus, previous studies investigating circadian rhythmicity of activity levels in patients with DOC might be subject to this bias. Furthermore, we found higher variability within the 24 h day (as indicated by higher intradaily variability [IV]) in the uncorrected data, thus suggesting a stronger fragmentation of the patients’ activity. In other words, IV increases when periods of low “real” patient activity are followed by strong activity initiated by moving the patient passively. Thus, while passive movements occur in a regular pattern *over several days* (i.e. resulting in more IS), the variability of the measured activity *within a day* is increased due to passive movements.

When looking at day-night variations of activity levels separately for patient groups, patterns between diagnoses stayed the same in the corrected and uncorrected dataset with MCS/EMCS patients showing stronger day-night effects than UWS patients (cf. *Figure 4*) as well as higher mean activity during day and night (cf. *Additional File 1: Figures S7 A-D*). Consequently, one might argue that the correction of actigraphy data is dispensable. However, as soon as the amount of passive movements differs between UWS and MCS/EMCS patients, we will get distorted results when contrasting actigraphy data between diagnoses. Even in our sample, where all of the patients were expected to receive equivalent levels of care, therapies and visits, the results from contrasting UWS and MCS/EMCS patients in the uncorrected data differed from the corrected data when looking at IV (cf. *Additional File 1: Figures S4 C-D*). Specifically, while UWS patients showed a significantly higher IV as compared to MCS/EMCS patients in the uncorrected dataset, no difference could be detected after correcting for passive movements.

Given the overestimation of circadian rhythms in the uncorrected dataset and the differing results of the two datasets when comparing diagnoses, we suggest using the corrected dataset when comparing actigraphy data of UWS and MCS/EMCS patients. Our analyses between diagnoses based on the corrected dataset revealed that the activity during both day and night was higher in MCS/EMCS patients than in UWS patients (cf. *Additional File 1: Figures S7 B+D*) and generally in patients with higher CRS-R scores (cf. *Additional File 1: Figure S9*). Also, MCS/EMCS patients had more pronounced circadian rhythms (i.e. normalized power; cf. *Figure 3B*). This indicates more preserved circadian rhythms in MCS/EMCS patients and is well in line with previous studies that investigated circadian rhythms in patients with DOC. Specifically, these studies showed that a higher integrity of circadian temperature and melatonin rhythms predict a richer behavioral repertoire, which is directly related to results of CRS-R assessments[5, 12]. Also on a brain level, day-night changes of EEG signal complexity are more pronounced in MCS than in UWS patients (with significantly higher signal complexity during day than during night[27]), and periods of “daytime wakefulness” and “night-time sleep” are better distinguishable in MCS than in UWS patients[28].

Besides this, the general usefulness of actigraphy data in severely brain-injured individuals especially for diagnostic and prognostic purposes seems questionable as the validity of motor data is severely limited by several factors such as motor impairments, spasticity and the usage of muscle relaxants in these patients. In a previous study of our lab, we did not find any relation between the IS of the patients’ physical activity levels and the CRS-R scores[5]. In the current study, IS correlated positively only with the motor subscale score, but not with the other subscale scores. Moreover, the effect was gone when contrasting UWS and MCS patients. We also did not find any significant correlations of the CRS-R scores with IV and the patient’s period length (i.e. deviation from the peak closest to 24 h). Therefore, we should be careful when drawing associations between circadian variations of physical activity in patients with DOC and consciousness levels (cf. *Additional File 1: Figure S9*). Instead, other measures such as hormones (i.e. melatonin(-sulfate)) seem to better describe circadian rhythms in patients with DOC; i.e. while we found a circadian rhythm in the corrected actigraphy data in only 13/30 patients (43%) in the current study (cf. *Additional File 1: Table S2*), 19/21 patients (90%) showed a circadian melatoninsulfate rhythm in our previous study[5].

## Conclusions

To summarize, our study shows that actigraphy from patients with DOC does not exclusively reflect the patients’ activity as it is strongly influenced by passive movements, which leads to an overestimation of the circadian rhythmicity of the activity initiated by the patients themselves. Consequently, actigraphy data needs to be corrected to allow for meaningful conclusions about circadian rhythms in patients with DOC. Considering this correction, we found that MCS/EMCS patients show higher mean activity during the day and night as well as stronger circadian rhythms than UWS patients. However, the general usefulness of actigraphy in patients with DOC should be considered carefully; especially with regards to frequent motor impairments, spasticity and the usage of muscle relaxants in these patients. Thus, while actigraphy is a tool that received increasing attention in measuring arousal because of its efficiency regarding costs and time, it must be treated with caution in clinical populations with severe motor impairments such as patients with DOC.

## Abbreviations

AI: anoxic-ischemic brain injury
CRS-R: Coma Recovery Scale – Revised
DOC: disorders of consciousness
EMCS: exit minimally conscious state
GCS: Glasgow Coma Scale
IQR: interquartile range
IS: interdaily stability
IV: intradaily variability
MCS: minimally conscious state
NTBI: non-traumatic brain injury
TBI: traumatic brain injury
UWS: unresponsive wakefulness syndrome

## Acknowledgements

We thank Sarah Haberl and Julius Köppen as well as the staff at the Albert-Schweitzer-Klinik Graz and the Gunther-Ladurner Pflegezentrum Salzburg for their constant support and help with the data collection.

## Declarations

### Availability of Data and Materials

The data that support the findings of this study are available from the corresponding author upon reasonable request.

### Authors’ Contributions

Study design: MaS, CB; Data acquisition: MA, CB, MR, GP, MoS, ABK; Data analysis and interpretation: MA, CB, MaS, MR; Drafting the manuscript: MA, CB, MaS; Critical revision of the manuscript: MA, CB, MR, GP, MoS, ABK, ET, MaS. All authors read and approved the final manuscript.

### Consent for publication

Not applicable.

### Ethics Approval and Consent to Participate

The study had been approved by the ethics commission of the medical university of Graz (21-506 ex 09/10) and the ethics commission of Salzburg (415-E/2034). Informed consent was obtained from the patients’ legal representatives.

## Additional Files

### Additional File 1

Supplementary Material (.docx)

- Additional information on **light levels, multiple CRS-R assessments** and **monitoring of passive movements**.
- **Figure S1.** Graphical user interface of the tablet in the patient room.
- **Table S1.** Circadian rhythm indices from the uncorrected actigraphy data.
- **Table S2.** Circadian rhythm indices from the corrected actigraphy data.
- **Figure S2.** Periodogram of the uncorrected (A) and corrected (B) actigraphy data from patient 26.
- **Figure S3.** Normalized power of the patients’ peaks closest to 24 h in uncorrected vs. corrected data.
- **Figure S4.** Interdaily stability (A+B), intradaily variability (C+D) and circadian rhythmicity (E+F) in MCS/EMCS vs. UWS patients separately for uncorrected and corrected data.
- **Figure S5.** Interdaily stability (A), intradaily variability (B), circadian rhythmicity (C) and circadian rhythm strength (D) in TBI vs. NTBI patients in the corrected data.
- **Figure S6.** Patients’ mean activity during day (7am – 9pm) vs. night (9pm – 7am) separately for uncorrected and corrected data.
- **Figure S7.** Mean activity levels during day (7am – 9pm) and night (9pm – 7am) in MCS/EMCS vs. UWS patients separately for uncorrected and corrected data.
- **Figure S8.** Mean activity levels during day (7am – 9pm) and night (9pm – 7am) in TBI vs. NTBI patients in the corrected data.
- **Figure S9.** Correlation Matrix.

