## Additional File 1 for "Actigraphy in brain-injured patients – A valid measurement for assessing circadian rhythms?"

SUPPLEMENTARY MATERIAL

M.Sc. Monika Angerer[1,2], Univ.-Prof. Dr. Manuel Schabus[1,2*], M.Sc. Marion Raml[1], M.D. Gerald Pichler[3], M.D. Alexander B. Kunz[4,5], M.D. Monika Scarpatetti[3], Univ.-Prof. M.D. Eugen Trinka[2,4], Dr. Christine Blume[2,6,7*]

*these authors contributed equally

**Affiliations:**

[1] University of Salzburg; Department of Psychology; Laboratory for Sleep, Cognition and Consciousness Research; Salzburg, Austria

[2] University of Salzburg; Centre for Cognitive Neuroscience Salzburg (CCNS); Salzburg, Austria

[3] Geriatric Health Centres of the City of Graz; Albert Schweitzer Clinic; Apallic Care Unit; Graz, Austria

[4] Department of Neurology; Paracelsus Medical University; Christian Doppler Medical Center; Salzburg, Austria

[5] Gunther Ladurner Nursing Home; Salzburg, Austria

[6] Centre for Chronobiology; Psychiatric Hospital of the University of Basel; Basel, Switzerland

[7] Transfaculty Research Platform Molecular and Cognitive Neurosciences; University of Basel; Basel, Switzerland

**Corresponding Author:**

Dr. Christine Blume
Centre for Chronobiology
Psychiatric Hospital of the University of Basel
Wilhelm-Klein-Str. 27
CH-4002 Basel

T: +41(0)61/325-5074

### Methods

#### *Light Levels*

Illuminance was kept <500 lux at eye level during the day (7 am – 9 pm) and <10 lux during the night (9 pm – 7 am) with a dim light (<10 lux at eye level) being switched on twice per night for 5-10 min for nursing activities. Light levels were monitored by continuous measurements with light sensors (wGT3X-BT Monitor, ActiGraph LLC., Pensacola, USA) and spot checks with a luxmeter (Dr. Meter, Digital Illuminance/Light Meter LX1330B). Relatives and nurses were instructed to cover the patients' eyes with sunglasses in case they went outside for a walk.

#### *Multiple CRS-R Assessments*

Multiple CRS-R assessments (i.e. 10 additional assessments) were obtained in 16 patients (8 females) on two days that were separated from the study week by a maximum of 22 days. On these two days, patients were assessed in regular time intervals spaced by 2.5 h around the time of the temperature maximum, which had previously been calculated from the skin temperature data obtained during the "study week". This was done because cognitive performance is expected to peak at the time around the temperature maximum[14, 29].

#### *Monitoring of Passive Movements*

To monitor passive movements and remove artifacts resulting from them, we recorded all events deemed relevant in the patient room using an application (https://github.com/wolli2710/HospitalTracker) that enabled clinical and research staff as well as visitors to indicate the type of activity that was performed by simply tapping the screen of a tablet in the patient room (e.g. nursing, patient in wheelchair, therapy, visit, etc.; cf. *Figure S1*).

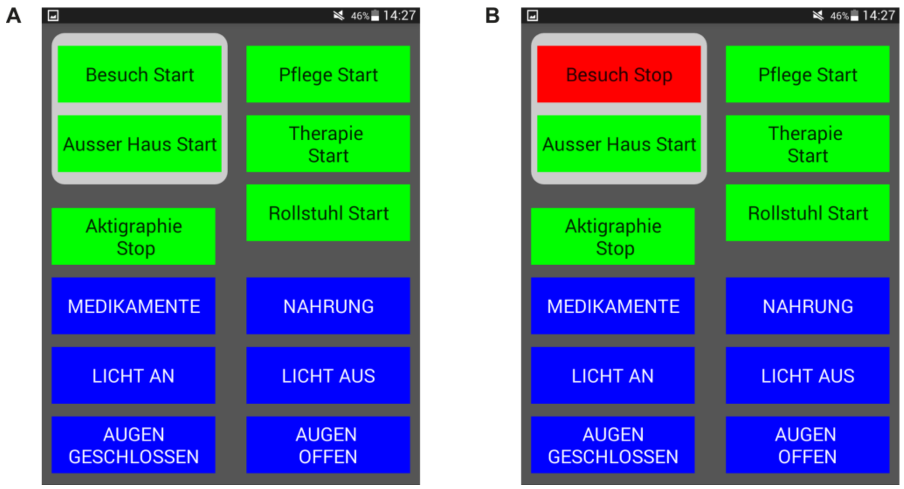


***Figure S1.* Graphical user interface of the tablet in the patient room.** While the green buttons have a start and end option (e.g. If "Besuch Start" is tapped **[A]**, the button turns red and changes to "Besuch Stop" **[B]**. It changes back to "Besuch Start" if it is tapped again.), the blue buttons are "single press buttons" (i.e. no start and stop option; only need to be pressed once at the time of occurrence). In green: Besuch Start = visit start, Ausser Haus Start = out of the building start, Aktigraphie Stop = actigraphy stop, Pflege Start = nursing start, Therapie Start = therapy start, Rollstuhl Start = wheelchair start. In blue: Medikamente = medication, Nahrung = nutrition, Licht an = lights on, Licht aus = lights off, Augen geschlossen = eyes closed, Augen offen = eyes opened.

### Results

#### *Individual Patient Results*

**Table S1.**
***Circadian rhythm indices from the uncorrected actigraphy data.***

| Patient ID | Mean day activity | Mean night activity | Interdaily Stability (IS) | Intradaily Variability (IV) | Peak Period | Normalized Power |
|---|---|---|---|---|---|---|
| P1 | 22.02 | 10.35 | 0.20 | 1.45 | 25.45 | 66.45 |
| P2 | 13.61 | 5.29 | 0.22 | 1.98 | 23.69 | 19.27 |
| P3 | 13.86 | 3.98 | 0.48 | 1.97 | 24.37 | 30.23 |
| P4 | 6.48 | 3.77 | 0.15 | 1.65 | 23.96 | 11.01 |
| P5 | 12.03 | 5.67 | 0.23 | 2.30 | 24.76 | 21.53 |
| P6 | 11.97 | 6.88 | 0.21 | 1.95 | 23.11 | 16.02 |
| P7 | 16.10 | 3.67 | 0.24 | 1.65 | 23.58 | 58.69 |
| P8 | 25.61 | 11.36 | 0.23 | 1.85 | 23.86 | 35.73 |
| P9 | 17.22 | 6.91 | 0.40 | 2.10 | 24.68 | 32.68 |
| P10 | 13.36 | 9.66 | 0.27 | 1.94 | 24.67 | 26.67 |
| P11 | 11.84 | 3.68 | 0.33 | 2.11 | 23.91 | 22.69 |
| P12 | 6.95 | 4.13 | 0.14 | 1.98 | 25.26 | 12.81 |

| | | | | | | |
|---|---|---|---|---|---|---|
| P13 | 20.21 | 7.94 | 0.29 | 1.94 | 22.03 | 28.24 |
| P14 | 11.66 | 5.93 | 0.14 | 1.37 | 23.20 | 44.75 |
| P15 | 6.65 | 2.82 | 0.55 | 2.21 | 23.87 | 14.08 |
| P16 | 160.84 | 30.97 | 0.25 | 0.81 | 23.86 | 542.81 |
| P17 | 15.85 | 4.39 | 0.40 | 2.12 | 23.80 | 44.22 |
| P18 | 6.08 | 2.80 | 0.15 | 1.50 | 24.17 | 18.67 |
| P19 | 518.87 | 77.06 | 0.31 | 0.72 | 24.33 | 516.47 |
| P20 | 5.07 | 2.10 | 0.28 | 2.14 | 24.59 | 14.29 |
| P21 | 305.43 | 132.43 | 0.49 | 1.00 | 24.29 | 319.41 |
| P22 | 10.39 | 4.30 | 0.18 | 2.01 | 24.02 | 24.58 |
| P23 | 22.22 | 15.13 | 0.17 | 1.71 | 22.61 | 12.98 |
| P24 | 14.49 | 4.99 | 0.26 | 1.85 | 24.55 | 39.76 |
| P25 | 27.26 | 13.74 | 0.35 | 1.35 | 24.82 | 58.25 |
| P26 | 9.66 | 1.91 | 0.30 | 2.12 | 24.21 | 45.46 |
| P27 | 17.89 | 8.84 | 0.17 | 2.01 | 22.35 | 16.41 |
| P28 | 13.94 | 2.91 | 0.20 | 1.80 | 23.94 | 61.61 |
| P29 | 128.82 | 215.35 | 0.08 | 0.38 | 23.77 | 136.00 |
| P30 | 13.32 | 4.75 | 0.34 | 1.99 | 23.52 | 30.05 |

Peak period and normalized power refer to the rhythm which is closest to 24 h. Markings in red indicate period lengths that deviate > 1 h from 24 h (i.e. 5/30 [17%] patients showed no circadian rhythm).

**Table S2.**
***Circadian rhythm indices from the corrected actigraphy data.***

| Patient ID | Mean day activity | Mean night activity | Interdaily Stability (IS) | Intradaily Variability (IV) | Peak Period | Normalized Power |
|---|---|---|---|---|---|---|
| P1 | 4.76 | 3.77 | 0.11 | 1.58 | 15.44 | 11.89 |
| P2 | 1.27 | 1.01 | 0.20 | 1.49 | 8.03 | 12.00 |
| P3 | 1.38 | 0.86 | 0.17 | 1.65 | 20.49 | 18.42 |
| P4 | 1.08 | 0.46 | 0.15 | 0.74 | 19.44 | 13.12 |
| P5 | 1.19 | 1.53 | 0.15 | 1.64 | 25.08 | 18.39 |
| P6 | 0.98 | 0.23 | 0.21 | 1.60 | 24.13 | 24.78 |
| P7 | 6.31 | 0.97 | 0.21 | 1.00 | 22.98 | 146.28 |
| P8 | 3.46 | 2.40 | 0.19 | 1.37 | 22.34 | 26.74 |
| P9 | 1.22 | 0.35 | 0.13 | 1.36 | 9.80 | 11.36 |
| P10 | 1.69 | 1.94 | 0.23 | 1.79 | 5.64 | 10.55 |
| P11 | 0.53 | 0.16 | 0.16 | 1.57 | 23.12 | 12.33 |
| P12 | 0.61 | 0.29 | 0.14 | 1.29 | 25.49 | 11.20 |
| P13 | 1.78 | 1.54 | 0.12 | 1.15 | 25.88 | 31.77 |
| P14 | 5.21 | 4.04 | 0.15 | 1.35 | 22.55 | 72.90 |
| P15 | 0.26 | 0.32 | 0.15 | 1.81 | 4.34 | 11.26 |
| P16 | 141.27 | 20.69 | 0.21 | 0.67 | 23.93 | 610.21 |
| P17 | 2.17 | 1.33 | 0.18 | 1.92 | 25.00 | 32.71 |
| P18 | 3.10 | 2.51 | 0.37 | 0.88 | 24.44 | 26.50 |
| P19 | 449.64 | 65.06 | 0.27 | 0.77 | 24.60 | 473.45 |

| | | | | | | |
|---|---|---|---|---|---|---|
| P20 | 1.14 | 0.51 | 0.13 | 1.22 | 25.11 | 31.82 |
| P21 | 242.42 | 124.82 | 0.37 | 0.97 | 24.17 | 215.98 |
| P22 | 0.63 | 1.96 | 0.23 | 1.49 | 25.37 | 18.25 |
| P23 | 13.20 | 6.93 | 0.17 | 1.90 | 23.08 | 18.70 |
| P24 | 0.47 | 0.77 | 0.15 | 1.82 | 24.22 | 10.48 |
| P25 | 11.71 | 8.92 | 0.20 | 0.49 | 24.10 | 138.44 |
| P26 | 0.02 | 0.00 | 0.16 | 1.16 | 24.56 | 15.20 |
| P27 | 5.20 | 4.68 | 0.33 | 1.13 | 25.66 | 41.35 |
| P28 | 0.87 | 0.54 | 0.18 | 1.40 | 24.25 | 11.31 |
| P29 | 123.74 | 210.51 | 0.08 | 0.41 | 23.62 | 131.61 |
| P30 | 3.44 | 0.87 | 0.13 | 0.99 | 22.23 | 23.88 |

Peak period and normalized power refer to the rhythm which is closest to 24 h. Markings in red indicate period lengths that deviate > 1 h from 24 h (i.e. 17/30 [57%] patients showed no circadian rhythm).

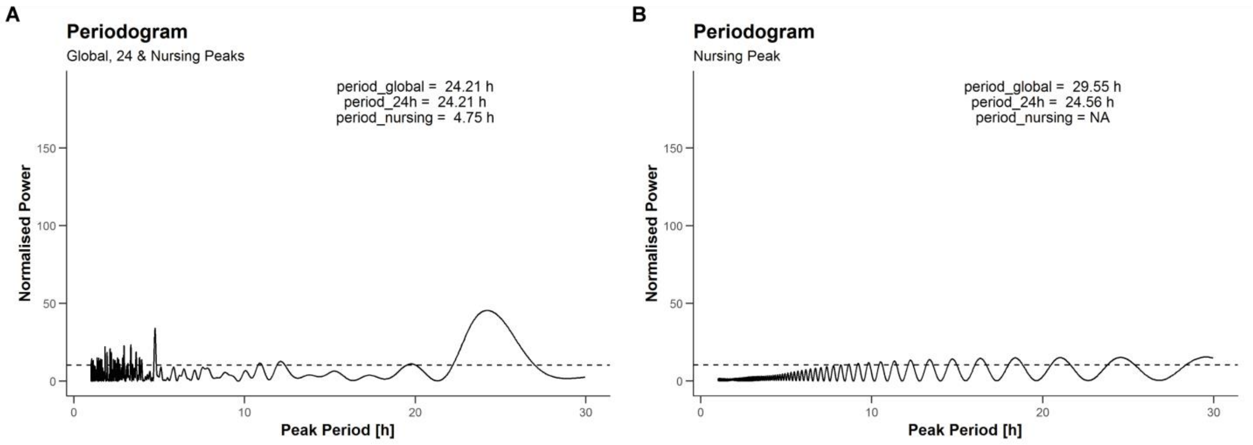


***Figure S2.* Periodogram of the uncorrected (A) and corrected (B) actigraphy data from patient 26.** Figure B shows that in extreme cases, no activity remains after correcting for passive movements. The dashed horizontal line indicates the significance level; with peaks exceeding this line being considered significant at $\alpha$ = 0.001. Normalized power describes the fit of a sine wave to the data. It is maximal where the sum of squares of the fitted sine wave to the data is minimal. Abbreviations: Period_global = significant peak with the highest normalized power, period_24h = significant peak closest to 24 h, period_nursing = significant peak with the highest normalized power between 3 and 5 h (period length).

*Circadian Rhythms*

**Uncorrected vs. Corrected**

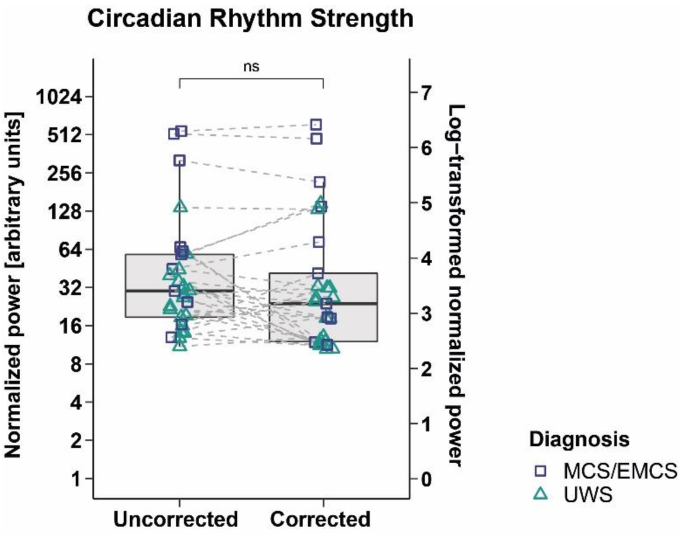


***Figure S3.*** **Normalized power of the patients' peaks closest to 24 h in uncorrected vs. corrected data.** The normalized power did not differ between uncorrected and corrected data. For better illustration, the data was log-transformed (right-hand y-axes); statistics were performed on the untransformed data (left-hand y-axes). Horizontal lines represent the medians, boxes the interquartile range (IQR; distance between the 1st [Q1] and 3rd quartile [Q3]), whiskers extend at most to Q1-1.5*IQR (lower whisker) and Q3+1.5*IQR (upper whisker). Abbreviations: ns = not significant, MCS = minimally conscious state, EMCS = Exit MCS, UWS = unresponsive wakefulness syndrome.

**MCS/EMCS vs. UWS**

The interdaily stability (IS) did not differ between diagnoses in both uncorrected ($Z(n_1$=11, $n_2$=18)=-0.47, *p*=.637, *r*=.09; cf. *Figure S4 A*) and corrected data ($Z(n_1$=11, $n_2$=18)=1.38, *p*=.169, *r*=.26; cf. *Figure S4 B*). While we found a significantly stronger intradaily variability (IV) in UWS as compared to MCS/EMCS patients in the uncorrected data ($Z(n_1$=11, $n_2$=18)=-2.20, *p*=.028, *r*=.41; cf. *Figure S4 C*), there was no significant difference in the corrected data ($Z(n_1$=11, $n_2$=18)=-1.42, *p*=.157, *r*=.26; cf. *Figure S4 D*). The deviation from 24 h did not differ between diagnoses in both uncorrected ($Z(n_1$=11, $n_2$=18)=0.52, *p*=.605, *r*=.09; cf. *Figure S4 E*) and corrected data ($Z(n_1$=11, $n_2$=18)=-1.53, *p*=.127, *r*=.28; cf. *Figure S4 F*). Furthermore, while MCS/EMCS patients showed a significantly higher normalized power of the peak closest to 24 h than UWS patients in the uncorrected data ($Z(n_1=11, n_2=18)=2.16$, $p=.031$, $r=.40$), this contrast was only visible by trend in the corrected data ($Z(n_1=11, n_2=18)=1.84$, $p=.065$, $r=.34$; cf. *Figure 3 B*).

To find out where the differences between datasets in IV and normalized power of the peak closest to 24 h come from, we ran comparisons between corrected and uncorrected data separately for diagnoses. Analyses revealed that UWS patients show significantly higher IV in the uncorrected data as compared to the corrected data ($Z(n=18)=-3.66$, $p<.001$, $r=.86$). In MCS/EMCS patients this differences was just available by trend ($Z(n=11)=-1.73$, $p=.083$, $r=.41$). When running the same analysis for normalized power, we did not find significant differences between the corrected and uncorrected data in UWS ($Z(n=18)=-1.22$, $p=.223$, $r=.29$) and MCS/EMCS ($Z(n=11)=-0.31$, $p=.756$, $r=.07$) patients.

This implies that – in our study sample – actigraphy data from UWS patients is more strongly influenced by passive movements than data from MCS/EMCS patients, which in turn leads to an overestimation of the IV in UWS patients in the uncorrected data. Thus, comparisons of actigraphy data between DOC diagnoses, which are not corrected for passive movements, can produce misleading results if UWS and MCS/EMCS patients receive a different amount of treatment.

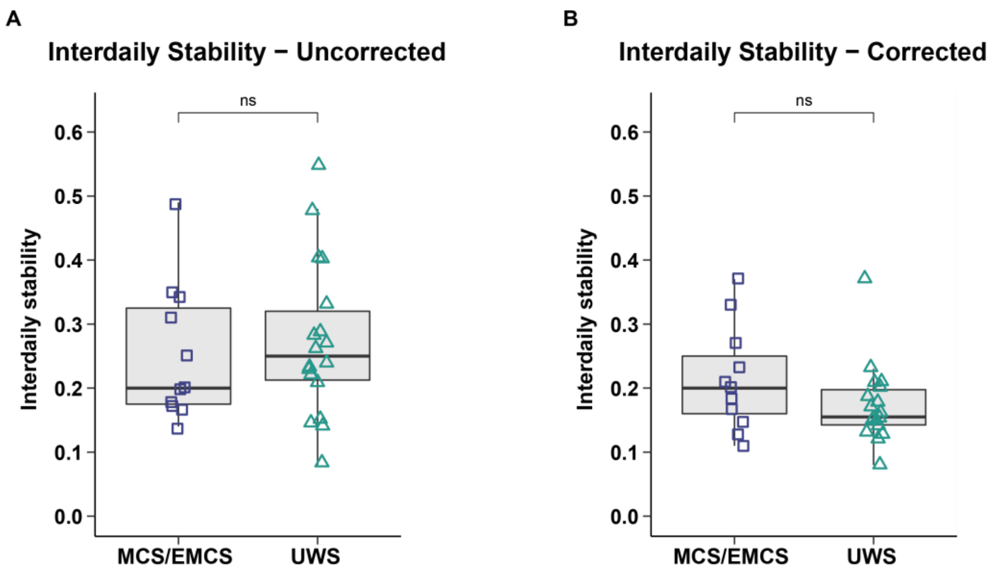

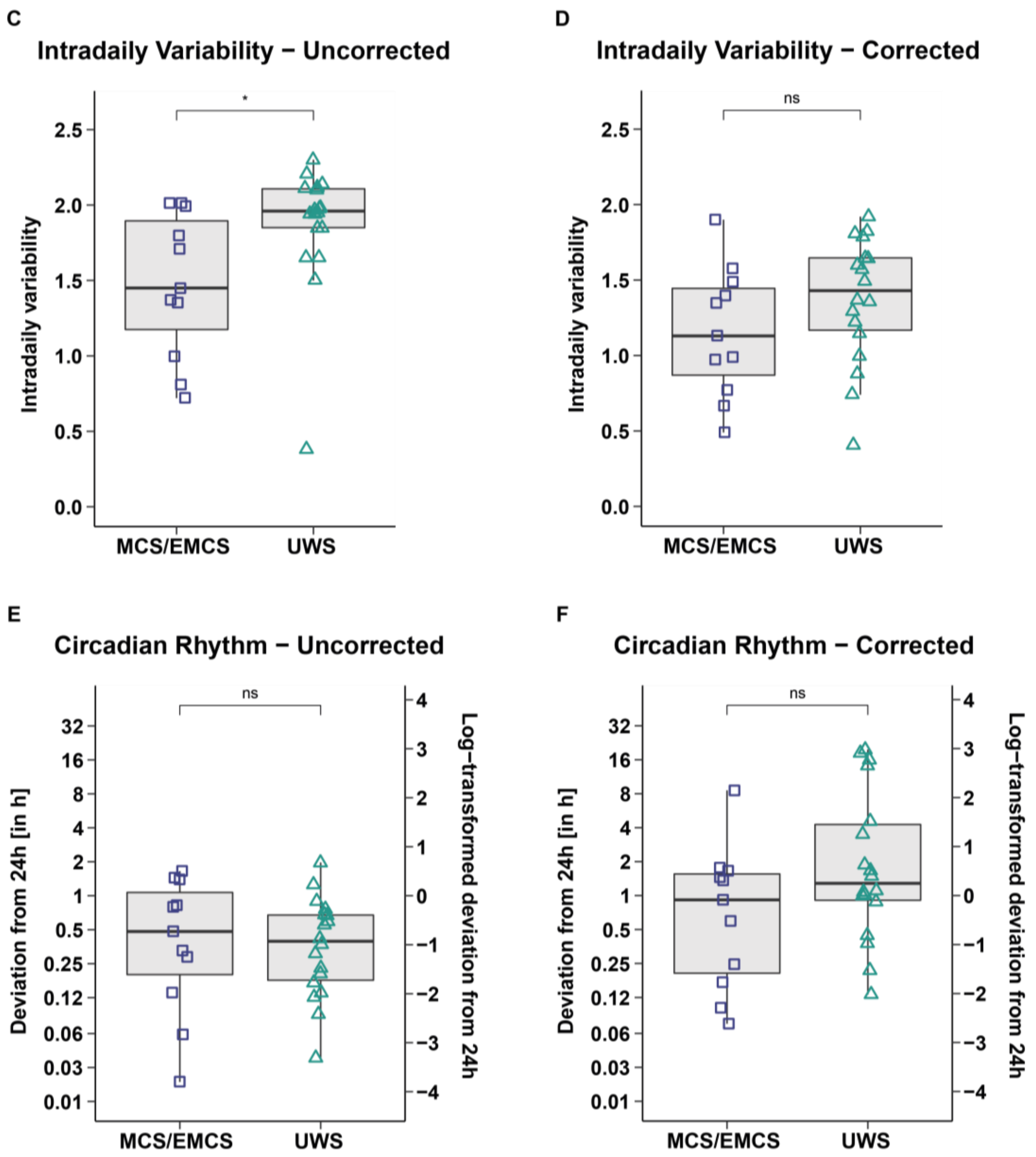


***Figure S4.* Interdaily stability (A+B), intradaily variability (C+D) and circadian rhythmicity (E+F) in MCS/EMCS vs. UWS patients separately for uncorrected and corrected data. A) Interdaily stability (IS) in the uncorrected data.** IS did not significantly differ between MCS/EMCS and UWS patients (IS approaches 0 for Gaussian noise and converges to 1 for perfect IS.). **B) IS in the corrected data.** IS did not significantly differ between MCS/EMCS and UWS patients. **C) Intradaily variability (IV) in the uncorrected data.** UWS patients showed significantly higher IV than MCS/EMCS patients (IV converges to 0 for a perfect sine wave [i.e. no IV] and approaches 2 for Gaussian noise. Values > 2 indicate an ultradian component with a period length of 2 h.). **D) IV in the corrected data.** IV did not significantly differ between MCS/EMCS and UWS patients. **E) Deviation of the patients' peak period from 24 h in the uncorrected data.** The deviation of the patients' peak period from 24 h did not differ between diagnoses. **F) Deviation of the patients' peak period from 24 h in the corrected data.** The deviation of the patients' peak period from 24 h did not differ between diagnoses. For better illustration of E+F the data was log-transformed (right-hand y-axes); statistics were performed on the untransformed data (left-hand y-axes). Horizontal lines represent the medians, boxes the interquartile range (IQR; distance between the 1st [Q1] and 3rd quartile [Q3]), whiskers extend at most to Q1-1.5*IQR (lower whisker) and Q3+1.5*IQR (upper whisker). Asterisks indicate significance: *$p \leq .05$, ns = not significant. Abbreviations: MCS = minimally conscious state, EMCS = Exit MCS, UWS = unresponsive wakefulness syndrome.

**TBI vs. NTBI**

We found no significant differences between etiologies (TBI vs. NTBI) in the IS ($Z(n_1$=10, $n_2$=19)=-1.11, *p*=.269, *r*=.21; cf. *Figure S5 A*), deviation from 24 h ($Z(n_1$=10, $n_2$=19)=0.96, *p*=.335, *r*=.18; cf. *Figure S5 C*), or normalized power ($Z(n_1$=10, $n_2$=19)=-1.15, *p*=.251, *r*=.21; cf. *Figure S5 D*) in the corrected dataset. However, patients with NTBI showed by trend a higher IV than patients with TBI ($Z(n_1$=10, $n_2$=19)=1.67, *p*=.094, *r*=.31; cf. *Figure S5 B*).

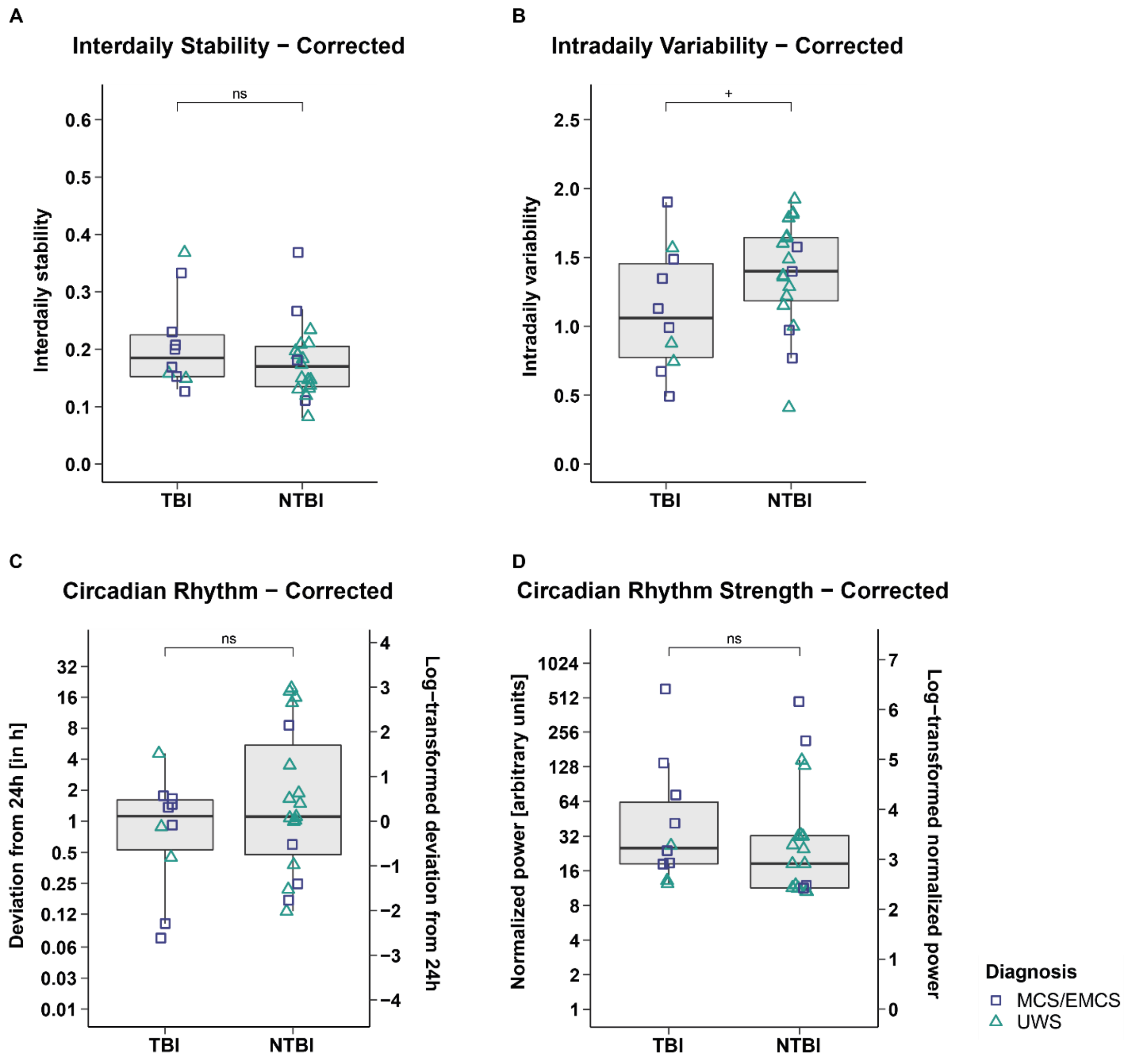


***Figure S5.*** **Interdaily stability (A), intradaily variability (B), circadian rhythmicity (C) and circadian rhythm strength (D) in TBI vs. NTBI patients in the corrected data. A) Interdaily stability (IS).** IS did not significantly differ between TBI and NTBI patients (IS approaches 0 for Gaussian noise and converges to 1 for perfect IS.). **B) Intradaily variability (IV).** NTBI patients showed by trend a higher IV than TBI patients (IV converges to 0 for a perfect sine wave [i.e. no IV] and approaches 2 for Gaussian noise. Values > 2 indicate an ultradian component with a period length of 2 h.). **C) Deviation of the patients' peak period from 24 h.** The deviation of the patients' peak period from 24 h did not significantly differ between etiologies. **D) Normalized power of the patients' peaks closest to 24h.** The normalized power did not significantly differ between etiologies. For better illustration of C+D the data was log-transformed (right-hand y-axes); statistics were performed on the untransformed data (left-hand y-axes). Horizontal lines represent the medians, boxes the interquartile range (IQR; distance between the 1st [Q1] and 3rd quartile [Q3]), whiskers extend at most to Q1-1.5*IQR (lower whisker) and Q3+1.5*IQR (upper whisker). Abbreviations: $^{+}p \leq .1$, ns = not significant, TBI = traumatic brain injury, NTBI = non-traumatic brain injury, MCS = minimally conscious state, EMCS = Exit MCS, UWS = unresponsive wakefulness syndrome.

### *Day vs. Night*

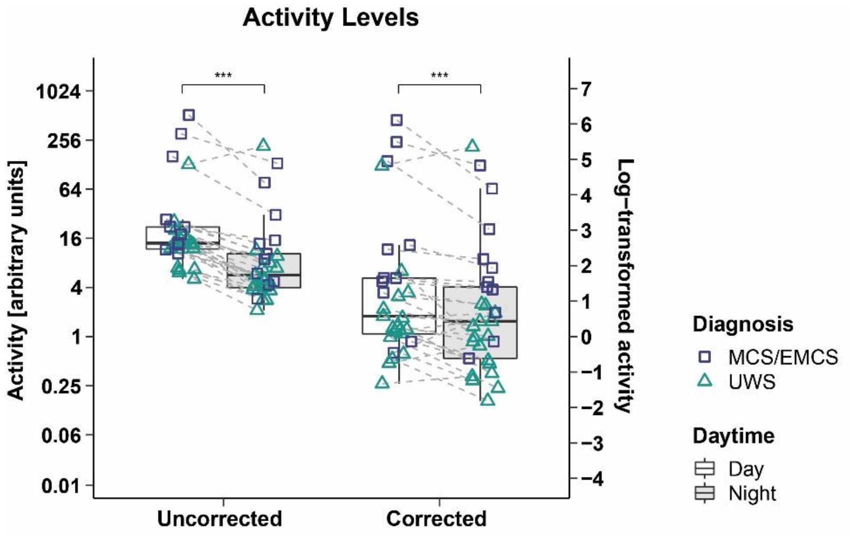


***Figure S6.*** **Patients' mean activity during day (7am – 9pm) vs. night (9pm – 7am) separately for uncorrected and corrected data.** The mean activity was significantly higher during day than during night in both the uncorrected and corrected dataset with stronger day-night effects in the uncorrected data. For better illustration, the data was log-transformed (right-hand y-axes); statistics were performed on the untransformed data (left-hand y-axes). Horizontal lines represent the medians, boxes the interquartile range (IQR; distance between the 1st [Q1] and 3rd quartile [Q3]), whiskers extend at most to Q1-1.5*IQR (lower whisker) and Q3+1.5*IQR (upper whisker). Asterisks indicate significance: $^{***}p \leq .001$. Abbreviations: MCS = minimally conscious state, EMCS = Exit MCS, UWS = unresponsive wakefulness syndrome.

### MCS/EMCS vs. UWS

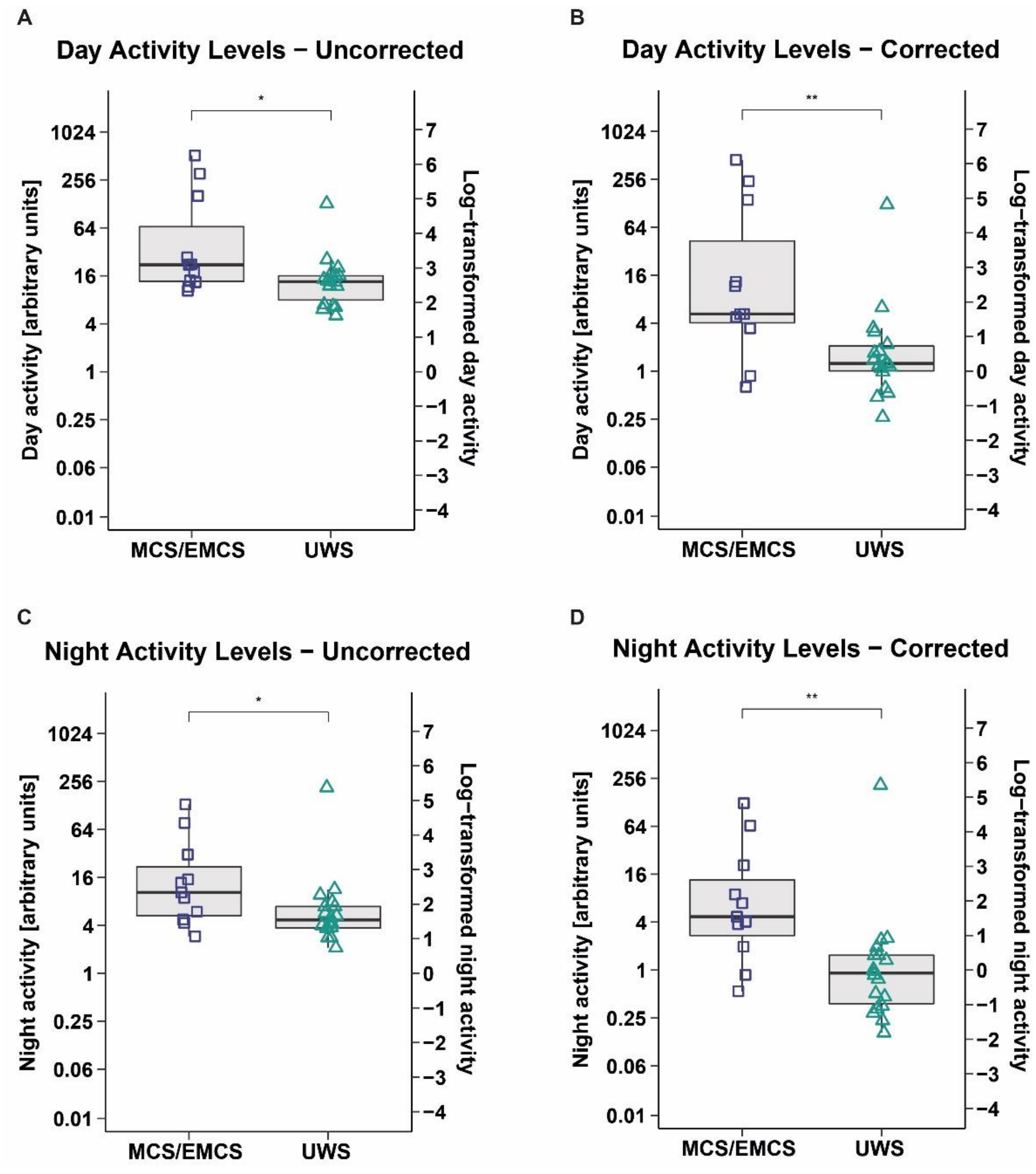


***Figure S7.*** **Mean activity levels during day (7am – 9pm) and night (9pm – 7am) in MCS/EMCS vs. UWS patients separately for uncorrected and corrected data. Day activity levels in the (A) uncorrected and (B) corrected data.** MCS/EMCS patients showed significantly higher mean activity during the day than UWS patients. **Night activity levels in the (C) uncorrected and (D) corrected data.** MCS/EMCS patients showed significantly higher mean activity during the night than UWS patients. For better illustration, the data was log-transformed (right-hand y-axes); statistics were performed on the untransformed data (left-hand y-axes). Horizontal lines represent the medians, boxes the interquartile range (IQR; distance between the 1st [Q1] and 3rd quartile [Q3]), whiskers extend at most to Q1-1.5*IQR (lower whisker) and Q3+1.5*IQR (upper whisker). Asterisks indicate significance: **$p \leq .01$, *$\leq .05$. Abbreviations: MCS = minimally conscious state, EMCS = Exit MCS, UWS = unresponsive wakefulness syndrome.

**TBI vs. NTBI**

We found no significant differences between patients with TBI and NTBI in the activity levels during day ($Z(n_1=10, n_2=19)=-0.92$, $p=.359$, $r=.17$; cf. *Figure S8 A*) and night ($Z(n_1=10, n_2=19)=-1.19$, $p=.233$, $r=.22$; cf. *Figure S8 B*) in the corrected dataset.

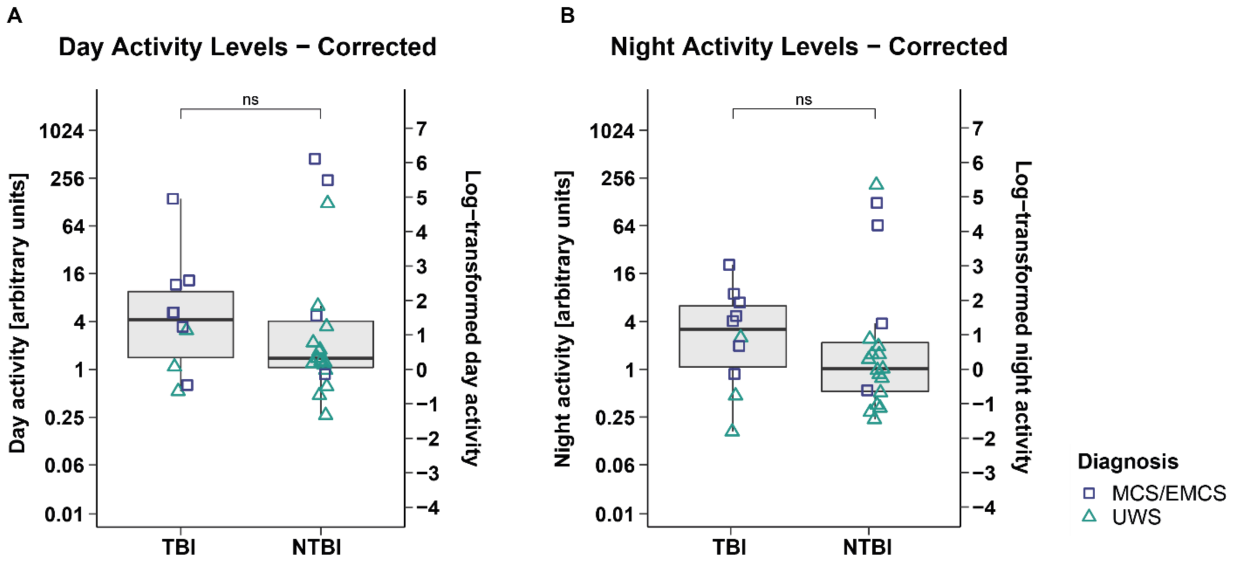


***Figure S8.* Mean activity levels during day (7am – 9pm) and night (9pm – 7am) in TBI vs. NTBI patients in the corrected data.** The mean activity during the day **(A)** and night **(B)** did not differ between etiologies. For better illustration, the data was log-transformed (right-hand y-axes); statistics were performed on the untransformed data (left-hand y-axes). Horizontal lines represent the medians, boxes the interquartile range (IQR; distance between the 1st [Q1] and 3rd quartile [Q3]), whiskers extend at most to Q1-1.5*IQR (lower whisker) and Q3+1.5*IQR (upper whisker). Abbreviations: ns = not significant, TBI = traumatic brain injury, NTBI = non-traumatic brain injury, MCS = minimally conscious state, EMCS = Exit MCS, UWS = unresponsive wakefulness syndrome.

*Correlation between CRS-R Scores and Actigraphy Data*

Our results indicate that the uncorrected data is not representative for measuring circadian rhythms in patients with disorders of consciousness as it is strongly influenced by passive movements. Therefore, we focused on the corrected data when correlating CRS-R scores with actigraphy data.

Results revealed positive correlations between CRS-R scores (sum score; auditory, visual, motor, and arousal subscale score) and mean activity during day (sum: $r_\tau(29)=.39$, $p=.004$; auditory: $r_\tau(29)=.35$, $p=.016$; visual: $r_\tau(29)=.32$, $p=.029$; motor: $r_\tau(29)=.38$, $p=.009$; arousal: $r_\tau(29)=.38$, $p=.011$) and night (sum: $r_\tau(29)=.42$, $p=.002$; auditory: $r_\tau(29)=.35$, $p=.014$; visual: $r_\tau(29)=.35$, $p=.019$; motor: $r_\tau(29)=.30$, $p=.037$; arousal: $r_\tau(29)=.39$, $p=.009$). Furthermore, a higher normalized power of the peak closest to 24 h was associated with a higher score on the motor subscale ($r_\tau(29)=.33$, $p=.025$) and by trend with a higher score on the visual subscale ($r_\tau(29)=.28$, $p=.059$). Patients' IS correlated positively with the motor subscale ($r_\tau(29)=.32$, $p=.031$). Please note that significant correlations between actigraphy data and the communication subscale are not reported because they cannot be interpreted due to less variance in the communication subscale scores (0 points: 24 patients, 1 point: 1 patient, 2 points: 4 patients).

When correlating actigraphy data with age and time since injury, we found that a higher age was associated a higher IV ($r_\tau(29)=.32$, $p=.016$) and by trend with less mean activity during day ($r_\tau(29)=-.23$, $p=.081$) (cf. *Figure S9*).

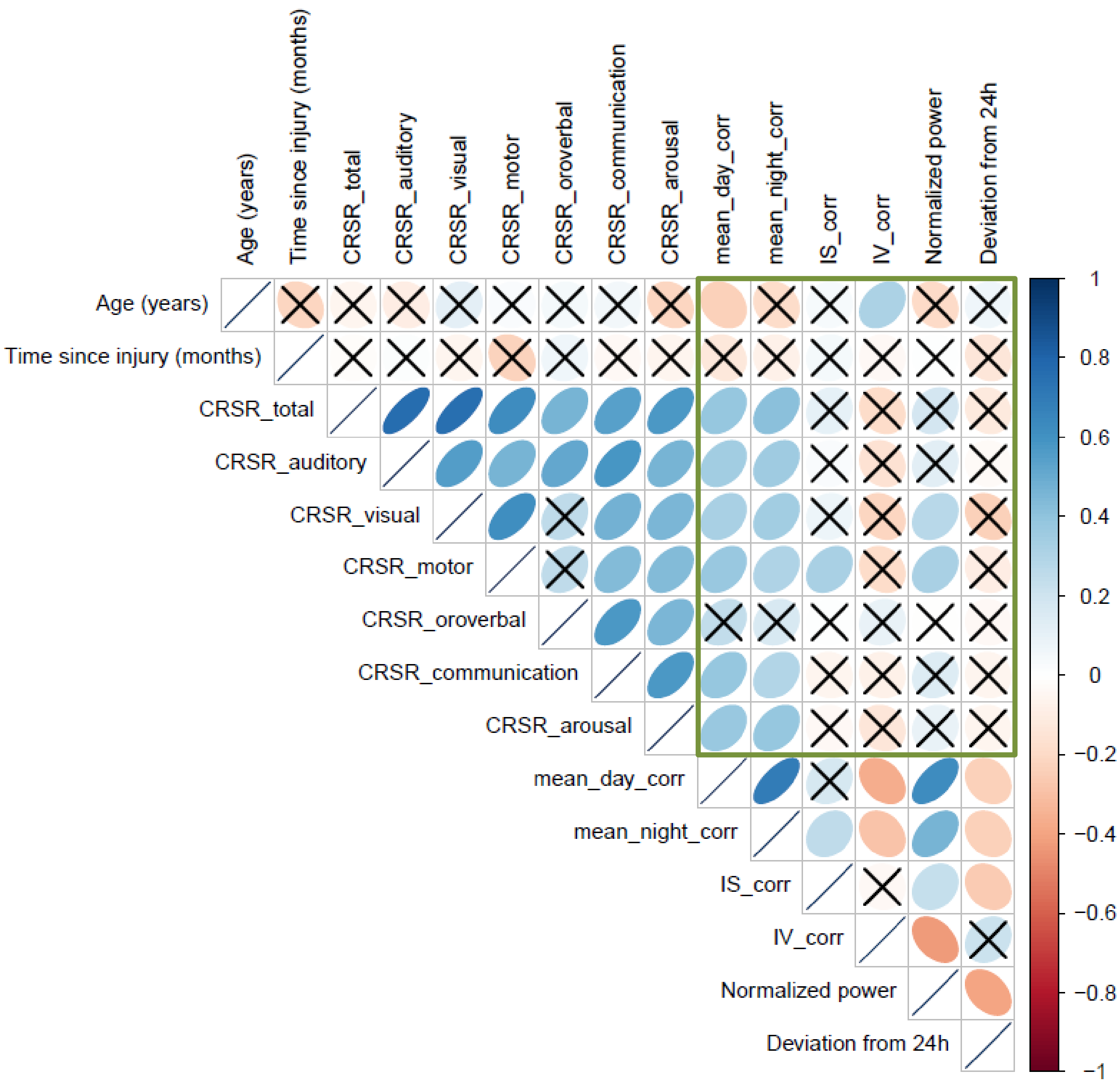


***Figure S9.*** **Correlation Matrix.** Positive correlations are displayed in blue and negative correlations in red. Color intensity and the size of the circle are proportional to the correlation coefficients (Kendall's Tau). Crosses indicate correlations with $p > 0.1$. Statistics for significant correlations in the green box are mentioned in the text above. Please note that significant correlations between actigraphy data and the communication subscale cannot be interpreted due to less variance in the communication subscale scores (0 points: 24 patients, 1 point: 1 patient, 2 points: 4 patients). IS = Interdaily Stability, IV = Intradaily Variability, corr = dataset corrected for passive movements.
